# Sensitive long-read amplicon sequence variant recovery with savont

**DOI:** 10.64898/2026.05.26.727271

**Authors:** Jim Shaw, Marie Riisgaard-Jensen, Kasper Skytte Andersen, Rasmus Kirkegaard, Morten Kam Dahl Dueholm, Heng Li

## Abstract

Long-read amplicon sequencing can profile longer sequences compared to short reads, but recovering amplicon sequence variants (ASVs) is a challenge for long, noisy reads. We present savont, an algorithm for recovering ASVs from modern long-read amplicons with mean accuracy ≥ 98%, including Oxford Nanopore Technologies (ONT) R10.4 and PacBio HiFi reads. Savont requires 5 to 16 times lower sequencing depth for full-length 16S rRNA ONT reads than previous methods and generates up to 95 times more ASVs for complex environments. Savont makes ASV-based analysis from shallow and noisy long-read amplicon sequencing feasible.

## 1 Introduction

Amplicon sequencing of conserved marker sequences has become a standard, cost-efficient analytical technique for diverse tasks [1,2]. Determining the exact marker sequences in a sample can lead to functional or taxonomic information, e.g., 16S rRNA genes for taxonomic profiling. A key computational task is to disentangle true variation from sequencing noise at single-nucleotide precision within these markers [3,4,5,6]; the inferred, noiseless sequences are called *amplicon sequence variants* (ASVs).

Traditional short-read amplicons target a few hundred base pairs, lacking the resolution of whole genome approaches [7]. Excitingly, long-read sequencing now enables longer amplicons, such as full-length 16S rRNA genes [8] or entire rRNA operons [9]. This increased length markedly increases the resolution, with applications to species or strain-level microbial profiling [10]. Key to accessing this resolution is generating true ASVs rather than operational taxonomic units (OTUs) [6] clustered at a fixed identity threshold (e.g., 97%). Thus, wet-lab approaches for ASVs [11,12] have been developed to circumvent the error rates of long reads. However, obtaining long-read ASVs from standard amplicon sequencing remains elusive.

The main methodological gap is that existing long-read ASV pipelines [13,14] must use “denoising” algorithms [4,5] designed for short reads. Denoising relies on identifying error-free reads within a set of reads. However, error-free long reads may be rare when amplicon lengths are long and sequencing noise is high (**Fig. 1a**). A recent study showed that simple mock communities with 8 species still require ≥ 50, 000 reads for full ASV recovery when using denoising algorithms with ONT R10.4 reads [14]. Thus, current methods may fail on low-depth samples or neglect low-abundance amplicons.

**Fig. 1:**
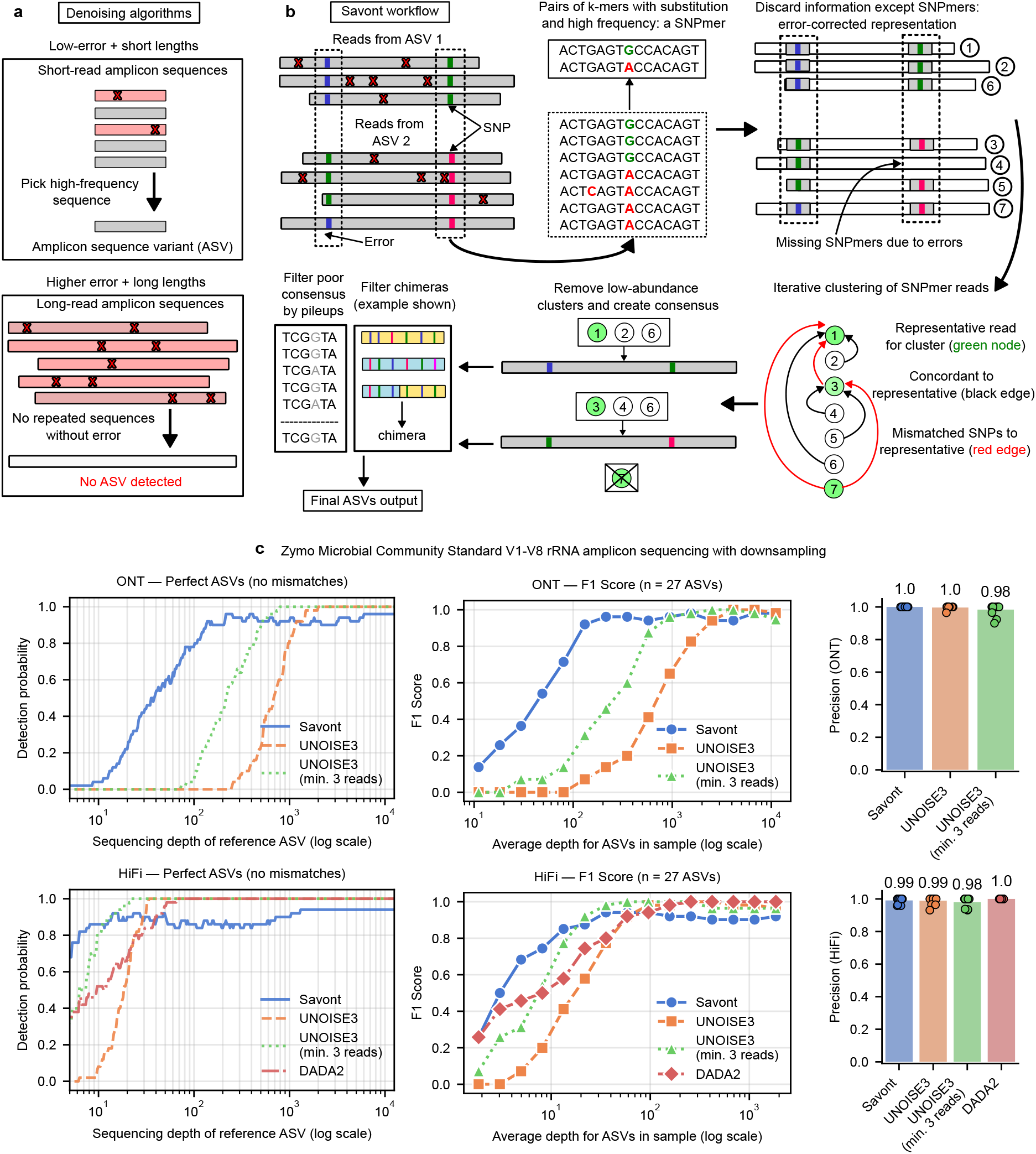
Savont recovers reliable ASVs through a cluster-and-consensus approach with polymorphic k-mers. **a**. Denoising algorithms assume the existence of error-free amplicons, which requires low error rates or shorter reads. Long-read amplicons violate these assumptions. **b**. Savont’s algorithmic workflow. **c**. Benchmarking on a V1-V8 16S rRNA Zymo amplicon dataset (D6305 / D6306) with 8 species and 27 true ASVs. Left: k-nearest neighbor regression (k = 50) across all 15 samples where the x-axis is the true ASV depth (determined by read mapping) and the y-axis is 1 if the ASV is recovered with 0 mismatches. Middle: F1 scores for recovered ASVs on the 15 downsamples. Average ASV depth (x-axis) is calculated as # of reads divided by 27. False positives are ASVs with ≥ 1 mismatch, and true positives are ASVs with 0 mismatches. Right: Mean precision (text) over all 15 datapoints.

In this work, we present savont, a new algorithm to generate ASVs from modern long-read amplicon sequencing with mean accuracy ≳ 98%, including ONT R10.4 reads and PacBio HiFi. Savont uses a cluster-and-consensus approach, which does not assume the existence of error-free reads. Through modern technologies [15] and algorithmic techniques [16], we demonstrate that it is possible to consistently recover low-frequency ASVs at single-nucleotide resolution. This leads to an order of magnitude higher sensitivity for ONT data than existing approaches, revealing hidden diversity in complex microbiome samples.

## 2 Results

### 2.1 Savont generates accurate ASVs at low depth on the Zymo mock community

At a high-level, savont clusters reads into putative ASVs and then generates an error-free consensus (**Fig. 1b**). The key challenge is to cluster erroneous reads that differ by even a single nucleotide polymorphism (SNP). We attack this problem by finding polymorphisms within the sample in a reference-free manner with *SNPmers* [16,17]: high-frequency pairs of k-mers that differ by a substitution in the middle base. Savont uses an efficient greedy clustering approach while ensuring that each read’s SNPmers within a cluster have the same middle base, thereby differentiating SNPs. Finally, consensuses with poor alignment pileups and putative chimericism [18] are removed (**Methods**).

We benchmarked savont against DADA2 [5] (for HiFi data) and UNOISE3 [4] (ONT and HiFi) on V1-V8 16S rRNA amplicons (≈ 1.5 kbp) and full rRNA operon amplicons (≈ 4 kbp). We used two sets of data sequenced from the Zymo Microbial Community Standard community: an ONT dataset from Riisgaard-Jensen et al [14] (D6305) and a HiFi dataset from Karst et al [11] (D6306). We discuss V1-V8 results in the main text (**Fig. 1c**) and show rRNA operon results in **Supplementary Fig. 1**. We ran all methods on 15 in silico downsampled communities to determine the impact of sequencing depth (runtime and memory usage in **Supplementary Fig. 2**). There are exactly 27 true ASVs in the V1-V8 dataset [14,12].

A critical question is how much sequencing depth is needed for generating ASVs. To answer this, we pooled all downsamples and performed k-nearest neighbor regression with ASV detection (1 = detected, 0 = not detected) as the response variable and true ASV coverage as the predictor (**Fig. 1c; left**). Savont recovers ASVs with 50% probability at 40x (ONT) and 5x (HiFi) depth. In contrast, UNOISE3 with default parameters requires 679x (ONT) and 19x (HiFi) depth for 50% detection probability, requiring 4-to 16-fold more reads. When lowering UNOISE3’s minimum error-free reads parameter from 8 (default) to 3, UNOISE3 requires 215x depth (ONT) and 7x depth (HiFi) for 50% detection. Thus, for HiFi, DADA2 (11x depth for 50% detection) and UNOISE3 were competitive in terms of sensitivity. A minor drawback of savont is the inability to differentiate a few intragenomic 16S rRNA gene copies at high depth, e.g., obtaining at most 26 / 27 ASVs in an ONT sample. Nevertheless, savont strongly outperforms existing methods for low-depth ONT sensitivity.

Framing perfect ASV detection as a classification problem (**Fig. 1c; middle and right**), savont’s improvement on low-depth communities was striking: for samples with mean ASV depth < 100x (i.e., < 27,000 reads), savont had a mean F1 score of 40.3% (ONT) and 74.8% (HiFi) compared to second-best F1 of 5.6% (ONT + UNOISE3 with minimum 3 reads) and 65.1% (HiFi + UNOISE3 with minimum 3 reads). Almost all ASVs were error-free: savont achieved a mean precision of 100% (ONT) and 99% (HiFi) across all 15 downsamples. Notably, savont output 0 false positives across all 15 ONT datasets compared to 10 for UNOISE3 with minimum 3 reads (**Supplementary Fig. 3**).

### 2.2 Savont unveils hidden ASV diversity on real environmental amplicon data

To evaluate savont on real samples, we ran all methods except DADA2 on four real ONT 16S rRNA (V1-V8) samples with > 1 million reads and in silico downsampling (**Fig. 2**) from Riisgaard-Jensen et al [14]. Savont outputs higher Shannon diversity and ASVs than UNOISE3 on all datasets, with a notable increase at low sequencing depth (**Fig. 2a**). In fact, UNOISE3 output no ASVs on 3/4 datasets when the number of reads was < 10, 000, and savont’s Shannon diversity was at least 2 points higher for all samples with ≤ 100, 000 reads. Savont’s sensitivity increase is especially important for complex microbiomes with many low-abundance ASVs: on the soil dataset, savont output 5639 ASVs compared to 378 (UNOISE3 minimum 3 reads) and 59 (UNOISE3 default).

**Fig. 2:**
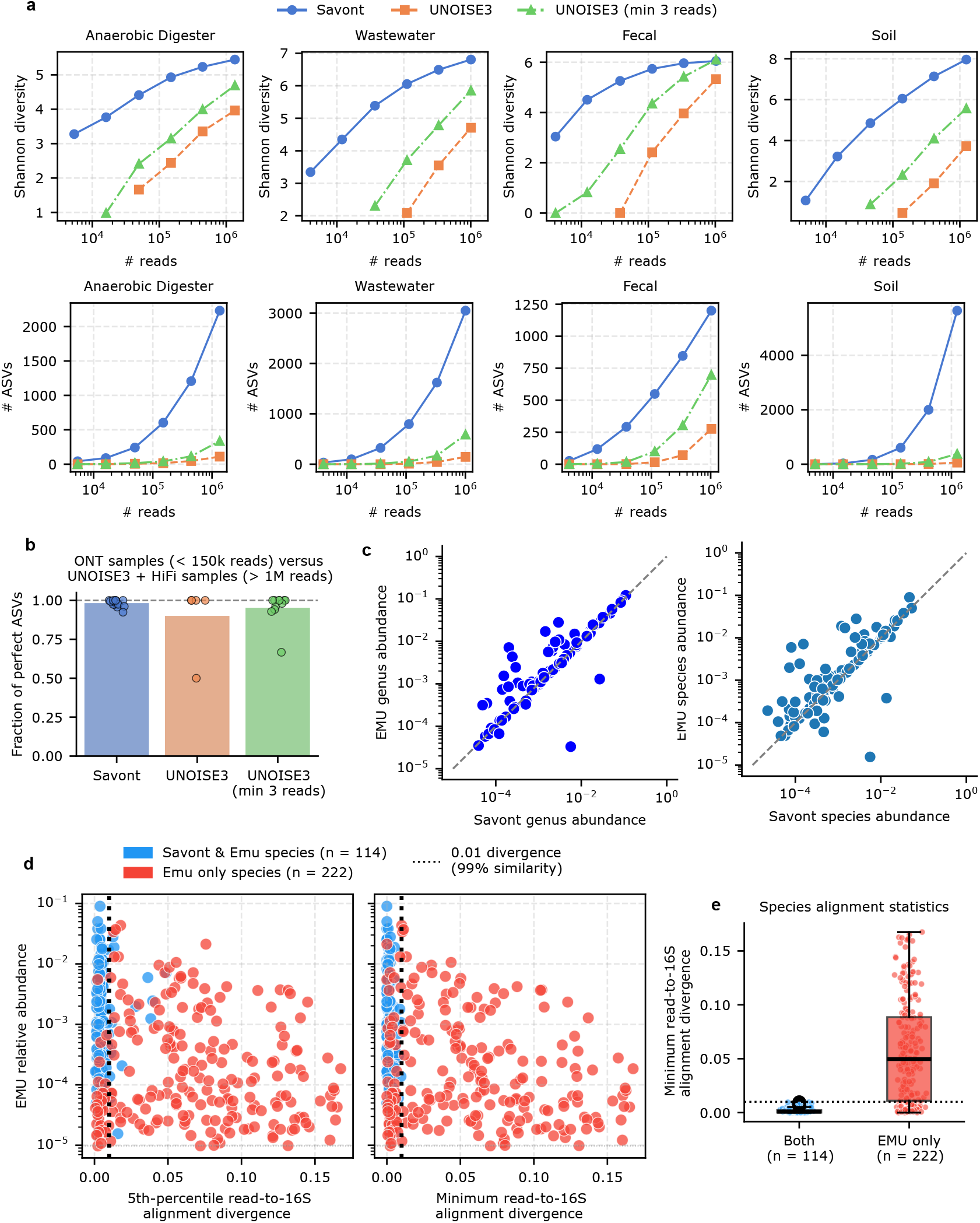
Results on real ONT 16S V1-V8 rRNA datasets. **a**. Top: Shannon diversity. Bottom: total number of ASVs recovered. **b**. Fraction of ASVs from downsampled ONT datasets (over all environments; *<* 150, 000 reads) mapping perfectly to a pseudo reference of ASVs recovered with UNOISE3 (minimum 3 reads) on the same samples but with *>* 1 million reads HiFi reads. **c**. Left: genus level abundances between savont and EMU on the fecal sample in (a). Right: species-level abundances. **d**. Alignment statistics for the ONT fecal 16S V1-V8 reads against EMU’s database. For each species, the 5th-percentile alignment divergence (left) and minimum alignment divergence (right) is calculated over all reads mapping against the species’ 16S rRNA sequences. **e**. Distribution of all species’ minimum 16S alignment divergence using the same data as panel (d). Boxplots show the median (line), 25th and 75th percentiles (box) and 1.5× the interquartile range (whiskers).

While it is difficult to infer false positive ASVs on real samples, the datasets of Riisgaard-Jensen et al. have high-depth HiFi sequencing as an orthogonal signal. We ran UNOISE3 (minimum of 3 reads) on the full HiFi samples, with each sample containing > 1 million reads, and used the results as a pseudo reference. We then compared the ONT ASVs from downsamples with < 150, 000 reads against the pseudo reference (**Fig. 2b**). On average, 98.2% of savont’s ASVs from the ONT subsamples (n=16 ONT downsamples) mapped perfectly (0 mismatches) against the pseudo reference. Thus, savont’s ASVs on real, complex samples have extremely high precision.

Lastly, we compared savont’s ability to generate ASV-based taxonomic profiles on the fecal sample to EMU [19], a read-mapping approach. We aligned savont’s ASVs to EMU’s database and defined species-level and genus-level hits to be > 99% [20] identity and > 94.5% [21] respectively; we note that savont’s ASVs could also be profiled via existing 16S classification methods [22]. For species and genera shared between EMU and savont, the Pearson R was 0.72 (species) and 0.96 (genus), showing broad concordance (**Fig. 2c**).

For this fecal sample, there was a large discrepancy in the number of species that EMU called (336) versus savont (115; 114/115 shared with EMU). For each species, we analyzed the minimum divergence across all minimap2 [23] alignments to its 16S genes (**Fig. 2d**). The 222 EMU-only species had a median minimum divergence of 6.1% compared to just 0.37% for the 114 species detected by both methods (**Fig. 2e**). Given that 1% divergence is a suggested species-level 16S rRNA gene threshold [20], this suggests that many EMU-only species calls represent overclassifications. False positive species calls for EMU also appear on two different ONT Zymo communities [24,14] (**Supplementary Fig. 4**). This issue has also been corroborated by Aja-Macaya et al. [25] in gut microbiomes. The latest EMU version allows users to set an alignment divergence threshold, which may aid in species-level precision but could also lead to misdetection of novel taxa. This shows the fundamental advantage of ASV-based profiling for resolving fine-scale ASV-to-database similarity.

## 3 Conclusion

We have described savont, a new computational method for resolving ASVs from long-read amplicon sequencing. Savont generates reliable ASVs from noisy, long amplicons of mean accuracy ≳ 98% without the need for UMIs or specialized sequencing approaches. Savont requires an order of magnitude less sequencing depth than previous computational methods through algorithmic innovations that leverage modern technologies. In conclusion, we believe that ASV-based profiling with noisy long-read amplicons on even shallow samples (< 100, 000 reads) is now reliable with savont.

## 4 Methods

The workflow of savont is outlined in **Fig. 1b**. Briefly, savont first generates pairs of polymorphic k-mers within a sample called SNPmers. SNPmers are essentially reference-free representations of bi-allelic SNPs that are anchored to a k-mer context. SNPmers are used to cluster the reads, enabling even single nucleotide polymorphisms to be distinguished. The clusters are turned into consensus sequences through partial order alignment [26,27], and these putative ASVs are filtered by removing low-quality sequences and chimeras.

### 4.1 Generating SNPmers

If a pair of k-mers within a sample differ only at the middle base, they are SNPmers. For example, if *AACAA* and *AAAAA* are both present in a sample, both *AACAA* and *AAAAA* are SNPmers. Due to sequencing errors, many k-mers within a sample will technically be SNPmers. However, our goal is to find SNPmers within the true ASVs. Thus, we will call SNPmers by filtering out sequencing errors using techniques analogous to variant calling methods [28].

Savont first counts all canonical k-mers (lexicographically smaller of forward and reverse k-mer; k = 17 by default) within a sample and removes k-mers that are not seen in both strands. Savont groups all k-mers into buckets sharing the same flanking *k* − 1 bases (but possibly different middle bases). The top two highest frequency k-mers within the bucket are considered candidate SNPmers and savont applies two tests: (1) a strand-bias test through Fisher’s exact test on the 2-by-2 contingency table of forward and reverse k-mer counts, and (2) a binomial test to filter out low-frequency k-mers; exact parameters are described in ref [16]. Pairs of k-mers passing these filters are retained.

A limitation of SNPmers is that they do not capture certain types of polymorphisms. For example, indels are not captured: if two sequences differ by a single indel, there are no SNPmers between them. In practice, indels are the primary error type of long reads [29], and indels are typically much harder than SNPs for variant calling [30]. Thus, we focus on SNPs in savont but leave indels for future work.

### 4.2 Indexing and filtering reads

Long reads are filtered by length and quality. Savont offers several preset length filters. By default, savont assumes full-length 16S rRNA amplicons and filters reads with length < 1100 or > 2000. Savont uses base qualities to filter out reads with mean estimated accuracy < 98%. The retained reads are indexed with the previously found SNPmers, but we require the middle SNPmer base have base quality > 25 by default.

In addition, k-mers (not necessarily SNPmers) are also sketched via FracMinHash [31] to retain approximately one out of *c* k-mers (*c* = 11 by default). We will use both the SNPmers and k-mers to do clustering in a two-step process.

### 4.3 Primary k-mer clustering

Savont performs two rounds of clustering. In the first round, all reads are quickly processed into primary clusters of roughly 95% identity. In the second round, savont processes each primary cluster separately. We use this hierarchical clustering process for two reasons. Firstly, it enables parallelization of the clustering step for computational efficiency. Secondly, highly divergent sequences may not have any shared SNPmers—SNPmer matching requires sufficient bases (i.e., the k-1 non-middle bases) be identical, which may be rare for divergent sequences. If two sequences share no SNPmers, then our proposed secondary clustering step will not work.

To do primary clustering, we use a greedy approach described at a high-level in Algorithm 1. We represent each cluster by a single representative read and a list assigning each read to a cluster. Savont iteratively queries reads against existing clusters and either assigns the query read to a cluster or creates a new cluster if it falls below the 95% similarity threshold to all cluster representatives. The similarity is quickly calculated by the standard k-mer identity estimator [32]

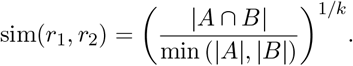

After clustering, we remove clusters with < 12 reads by default (reduced to 4 with `--hifi`). We found that a minimum of 12 reads (ONT) or 4 reads (HiFi) removes many false positive clusters and is sufficient for error-free ASV consensuses.

To perform the FindBestMatch step in Algorithm 1, we use a hash table index with a MinHash approach [33,34]. Because marker sequences can be highly conserved, a single k-mer (default k=17) can be shared between many sequences. Therefore, we hash all k-mers, take the bottom 3 k-mers (subject to the hash ordering) as a signature, and use this signature as a key. The 3-tuple is more discriminative than a single k-mer and thus more efficient for querying. We use 20 different hash functions, giving 20 signatures per read. To find the best match, we simply query a read’s signatures against the hash table to find similar reads, and then savont applies sim(*r*_1_, *r*_2_) against all candidate reads.

#### Algorithm 1

Greedy Primary K-mer Clustering

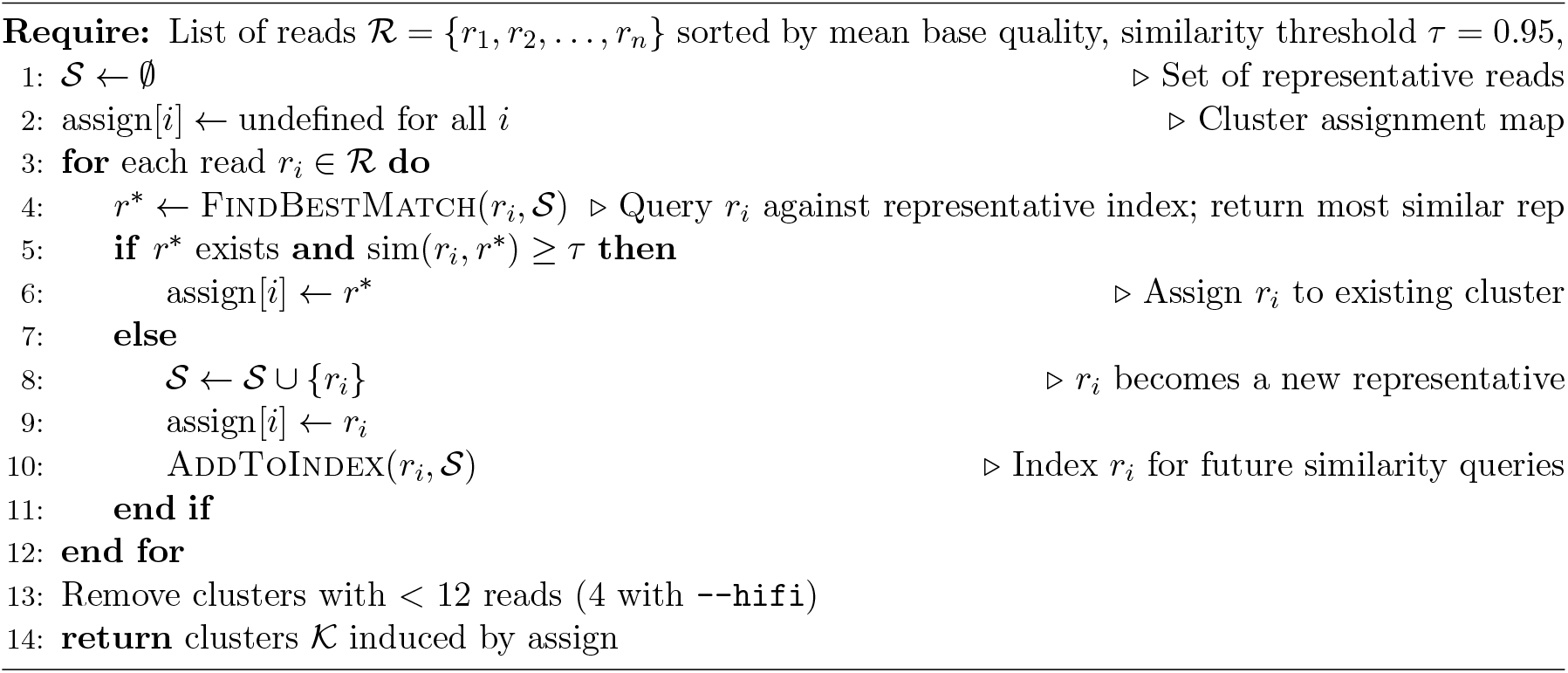

### 4.4 Secondary SNPmer clustering

In the secondary clustering step, primary clusters are refined in parallel through a greedy algorithm (Algorithm 2). The idea is broadly similar to the primary clustering algorithm. The key difference is that instead of a similarity threshold, for two reads to be in the same cluster their SNPmers must be concordant: no two SNPmers can have differing middle bases.

#### Definition 1 (Concordance).

*Given two partner SNPmers with identical flanking bases, for example AACAA and AAAAA, they are discordant if the middle base differs and concordant otherwise. Similarly, two reads are concordant if all SNPmer pairs are concordant. Otherwise, they are discordant*.

Findconcordantindexed uses an index to find concordant reads as follows. When ≤ 1000 clusters, we use a SNPmer hash table index: keys are SNPmers with their middle bases masked, and the values are the cluster representatives. Savont then queries a read’s masked SNPmers against this index and then compares the middle bases. If any representative is concordant to the query, then savont adds the query to the concordant representative’s cluster.

When sequencing depth is high, we found that many noisy clusters can form—even a single erroneous substitution may give rise to a noisy cluster. Therefore, if > 1000 clusters, we proceed through FindConcordantIterative instead. This checks the query against all representatives iteratively and stops after a concordant representative is found. This is faster in practice because most reads are concordant to an ASV representative early on in the iterative process, so only a few comparisons are needed for most reads.

### 4.5 Reclustering and refinement

The SNPmer clusters from Algorithm 2 are subject to a final reclustering step outlined in Algorithm 3. The previous clustering algorithms rely on a single representative read per cluster, but representatives can have errors, leading to missing SNPmers and poor initial clustering. Therefore, the reclustering step proceeds by building consensuses of SNPmers for each cluster and redistributing reads using the SNPmer consensuses, which are more reliable than just a single representative.

#### Algorithm 2

Greedy SNPmer Clustering

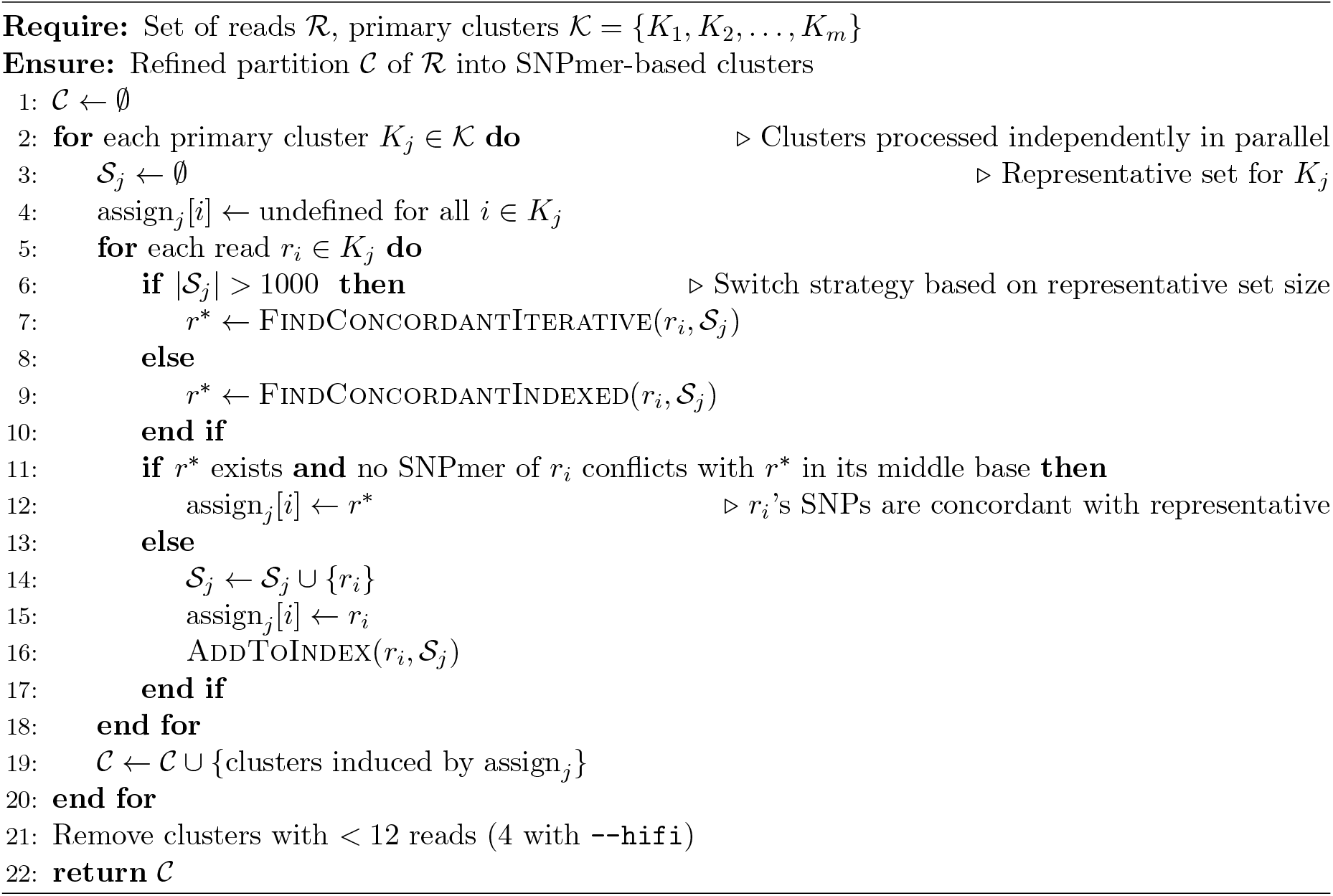

#### Algorithm 3

Iterative SNPmer-based reclustering

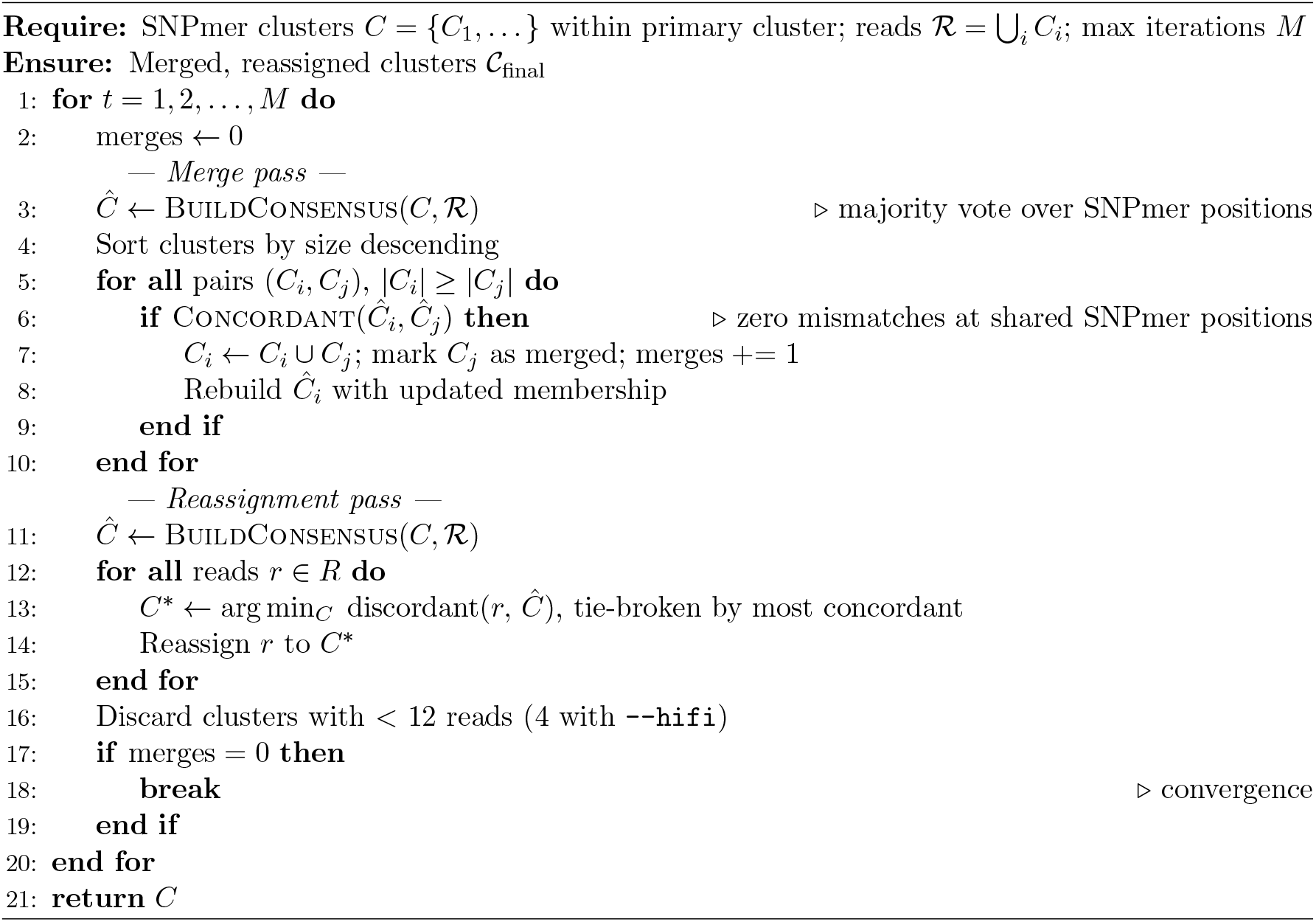

In Buildconsensus, for each cluster *C*_*i*_ we build a SNPmer consensus by removing all SNPmers appearing < 1/6 × |*C*_*i*_| times. Then, we iterate through all discordant SNPmer pairs and retain only the most abundant SNPmer to obtain a SNPmer consensus.

We then merge all cluster pairs (*C*_*i*_, *C*_*j*_) that are concordant: two cluster consensuses are concordant if all SNPmers are concordant and > 97.5% of *C*_*i*_ or *C*_*j*_’s consensus SNPmers are present in the other consensus. This last condition is to ensure we don’t merge consensuses that are very different (and thus don’t share SNPmers).

In the reassignment step, we query each read against all SNPmer consensuses and assign it to a consensus with minimal discordant SNPmers. In the case of tie breaks, we assign the read to the consensus with the maximal concordant SNPmers. Finally, we repeat this entire procedure until stability (i.e., no more merges) or we reach *M* = 10 iterations.

### 4.6 Consensus generation

For each cluster 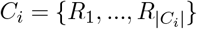 from Algorithm 3, we use SIMD Partial Order Alignment (SPOA) [27] to generate a consensus sequence (i.e., an ASV). We use *M* = min(75, |*C*_*i*_|) reads for SPOA for computational efficiency purposes. We then align all *M* reads back to the resulting consensus (using minimap2-rs [23]) and remove low-quality consensus ends with depth < 1/4 × *M*.

### 4.7 Empirical quality calibration and pileup filtering

Low-quality consensuses from the previous step may exist for a variety of reasons, for example, poor clustering or systematic errors within reads. The main idea of savont is to gauge the quality of a consensus by aligning the cluster’s reads back and examining alignment statistics. Low-quality consensuses will have ambiguous bases, and savont will filter them out.

For each cluster *C*, we take a subset *M* ^*′*^ = min(250, |*C*|) of its reads and align them back to the consensus. We then take the top 10% largest clusters, and for all positions with < 5% alignment error, we let

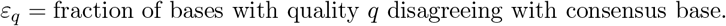

*ε*_*q*_ are the calibrated error rates given base quality *q*.

After calibration, for each position *p* in a consensus string, we define a pileup base quality under a simple probabilistic model as follows. Let a pileup at *p* be represented *A*_*p*_ = {(*b*_1_, *q*_1_), …, (*b*_*n*_, *q*_*n*_)} where *b*_*j*_ are aligned nucleotides or indels, and *q*_*j*_ are base qualities. We define insertions and deletions to be a special character with fixed *q* = 48, a heuristic to approximate ONT R10.4 indel rates. Given a reference base *b*, define

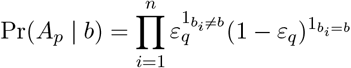

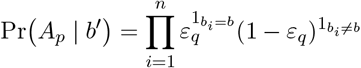

where 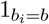 is 1 if *b*_*i*_ = *b* and 0 otherwise. Pr(*A*_*p*_| *b*) models the probability of observing the pileup given independent alignments with base qualities, and the *b*^*′*^ indicates “not *b*”. Finally, let *b* be the consensus base and define the pileup alignment score to be

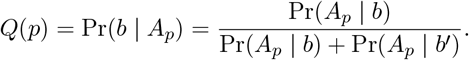

Here, we use Bayes’ theorem with uniform priors. Finally, we filter out a consensus if any position *p* within the consensus has ln(1 − *Q*(*p*)) > −*T*, where *T* = 30 is a default threshold.

### 4.8 Merging similar sequences

True low abundance ASVs and erroneous ASVs are difficult to distinguish based on abundance alone. A recurring sequencing error for reads from a highly abundant ASV can lead to a moderately abundant but erroneous ASV. A key insight is that while erroneous ASVs do not have low abundance in general, they have low abundance relative to a similar, high abundance ASV [4,35].

To remove erroneous consensuses, we follow the idea of UNOISE. Define *S*_*i*_ to be the ASV sequence from cluster *C*_*i*_. We align all pairwise consensuses with minimap2-rs and filter consensus *S*_*i*_ if there exists consensus *S*_*j*_ such that

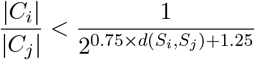

where *d*(*S*_*i*_, *S*_*j*_) is edit distance. The intuition is that the higher the edit distance, the less likely it is to be an erroneous ASV, thus requiring less strict abundance filtering.

### 4.9 Filtering chimeras

The final filtering step is removing PCR chimeras. The main idea is to see if a consensus can be decomposed as two higher abundance consensuses joined together. Thus, we proceed by computing pairwise sequence identities between all consensus sequences using minimap2-rs, restricting comparisons to pairs where one sequence has at least 3-fold greater read depth than the other.

A consensus *S* is flagged as a chimera and removed under the following condition: its left portion matches a > 3× higher-depth parent perfectly, and its right portion matches a distinct > 3× higher-depth parent perfectly, with the two matching segments together spanning at least max(0.8, 0.9 · s_*LR*_) of *S*, where *s*_*LR*_ is the identity between the two parents. The two parents must also be sufficiently divergent (identity < 0.97, or < 0.995 if both have at least 10× the depth of *S*).

### 4.10 Final ASV quantification

After generating consensus sequences, these ASV abundances are refined by re-mapping all reads to this final ASV set and using an expectation-maximization (EM) algorithm. This step resolves multi-mapping reads that arise from highly similar ASVs sharing polymorphic marker sequences.

Each read is first compared against all ASVs using their SNPmer. Candidate ASVs are scored by the following ratio:

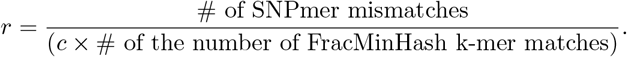

Here, *c* (default = 11) is the FracMinHash downsampling ratio. Under a random substitution model of sequence evolution, *r* is a statistical estimator for sequence divergence [16]. ASVs passing a threshold *r* < 0.005 are kept. Among passing candidates, savont proceeds with full pairwise alignment using minimap2-rs. Reads mapping with equal number of mismatches to multiple ASVs are grouped into equivalence classes [36,37], each representing a set of ASV targets that cannot be distinguished by that read.

Savont runs the EM algorithm on the equivalence class counts under a standard multi-mapping probability model to infer the true abundances [38]. Assuming *N* ASVs are output, we initialize ASV abundances at step 0 as

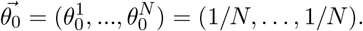

The output of the mapping stage is a set of equivalence classes *E* = {*e*_1_, …,} where each equivalence class is a set of ASVs *e* = {*ASV*_*j*_, …} with count *c*_*e*_. Then the EM update step can be written as [38]

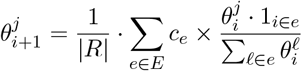

where 1_*i*∈*e*_ is a 1-0 indicator variable if *ASV*_*i*_ is in *e*. We run this procedure until the maximum per-ASV abundance change falls below 0.01/|*R*| or 10,000 iterations are reached, and then output the final 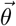.

### 4.11 Database classification and taxonomic profiling

Savont’s resulting ASVs have depth information and can be used by standalone ASV classifiers and taxonomic profilers, for example homology search methods (e.g., BLAST) [39] or k-mer-based methods [22,40]. To facilitate practitioner use, we have reimplemented two classification methods in savont: a minimap2-based homology search method and Edgar’s sintax algorithm [40]. For the homology-based method, ASVs with > 99% identity and > 94.5% identity against a database are classified as species and genus-level hits respectively; analogous thresholds for higher taxonomic ranks were taken from Yarza et al [20,21]. Savont currently supports downloading and analysis of the SILVA non-redundant 99% SSU database [41], EMU’s database [19], and Greengenes2 [42] for both classification methods.

### 4.12 Benchmarking setup

ONT R10.4 (with super-high accuracy basecalling) Zymo Microbial Community Standard reads were taken from Riisgaard-Jensen et al. [14] (specifically, technical replicate #1), who generated true ASVs using a UMI-based method [11]. HiFi rRNA operon reads were taken from Karst et al. [11]. Notably, Lin et al. [12] found substantial low-frequency contamination in the Karst et al. data. Therefore, if an ASV from the Karst et al. dataset had > 10 mismatches to our true ASVs, we did not consider it a false positive. We found that none of the ASVs with > 10 mismatches had > 500 bases perfectly aligning to another ASV, indicating they were unlikely to be chimeras.

Raw reads were first adapter-trimmed with Porechop, then primer-trimmed and filtered with cutadapt [43] v5.2 with parameters

`cutadapt -g AGRGTTYGATYMTGGCTCAG…TGYACWCACCGCCCGTC --rc --discard-untrimmed -e 0.1`

In the case of full 16S rRNA operon reads (**Supplementary Fig. 1**), we used the primers AGRGTTYGATYMTGGCTCAG…GTTTGGCACCTCGATGTCG instead.

To assess performance across sequencing depths, trimmed reads were subsampled to multiple fractions using seqtk v1.4 (https://github.com/lh3/seqtk). For the ONT reads, we subsampled reads between 300 to 300,000 reads with n=15 datapoints evenly spaced in log-scale. For HiFi, we used the same procedure but between 50 to 50,000 reads. ASVs were generated with the following commands:

1. Savont v0.4.0 (`savont asv`; default parameters and `--hifi` when appropriate.)
2. UNOISE3 v12.0 (<preformate>usearch -fastx uniques -relabel Uniq -minuniquesize 2 -fastaout uniques.fa; usearch -unoise3 uniques.fa -zotus asvs.fa</preformate>)
3. UNOISE3 v12.0 except with`usearch -unoise3 uniques.fa -zotus asvs.fa -minsize 3`
4. DADA2 via QIIME2 [44] version qiime2-amplicon-2024.10 with `qiime dada2 denoise-ccs`

All tools were run with 20 threads if possible on a Intel(R) Xeon(R) Silver 4316 CPU @ 2.30GHz machine. The resulting ASV sequences were aligned to references using minimap2. EMU v3.6.2 was run with default parameters and default database (https://osf.io/56uf7 version March 13, 2023).

Reference ASV coverages at each subsampling depth were independently quantified by aligning reads with minimap2 and applying samtools for depth estimation. Wall-clock runtime and peak resident memory for each denoising step were recorded using Snakemake’s [45] built-in benchmarking. For UNOISE3 and UNOISE3-min3, the dereplication and denoising runtimes were summed and peak memory was taken as the maximum across both steps. Claude Sonnet v4.5 and v4.6 via Claude Code were used to assist in writing the benchmarking scripts with downstream manual inspection of the scripts.

### 4.13 Analysis of complex communities

Savont and UNOISE3 were run on the four complex communities from Riisgaard-Jensen et al’s V1-V8 ONT R10.4 amplicon datasets. Only the first technical replicate was used for the ONT samples. To generate the PacBio HiFi pseudo reference (**Fig. 2b**), all four technical replicates from Riisgaard-Jensen et al. were concatenated. Given a set of ASVs with abundances *A*_1_, …, *A*_*n*_, Shannon diversity was calculated as 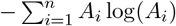. We determined ASV abundances for UNOISE using the `usearch –otutab` command and normalizing. For the analysis of EMU’s alignments (**Fig. 2d**), we used `emu --keep-files` to retain the SAM file from EMU.

## 5. Data availability

The Riisgaard-Jensen et al. dataset ONT V1-V8 and rRNA operon datasets are available in the Sequence Read Archive at PRJNA1389832. The Karst et al. HiFi Zymo Microbial Community Standard (D6306) dataset is available at ERR3813246. The Zymo D6305 dataset from ONT is available at https://epi2me.nanoporetech.com/zymo_16s_2025.09/.

## 6. Code availability

Savont is open source under the MIT license. Documentation and code are available at https://github.com/bluenote-1577/savont. Benchmarking scripts are available at https://github.com/bluenote-1577/benchmarks_savont.

## 7 Acknowledgments

This work is supported by US National Institute of Health grant no. R01HG010040 to H.L. J.S. is supported by a Natural Sciences and Engineering Research Council of Canada (NSERC) Postdoctoral Fellowship award (no. PDF-587396). M.R.-J. and M.K.D.D. were funded by the Novo Nordisk Foundation (REThiNk, grant NNF22OC0071498, M.K.D.D)

## A Supplementary Materials

**Supplementary Figure 1:**
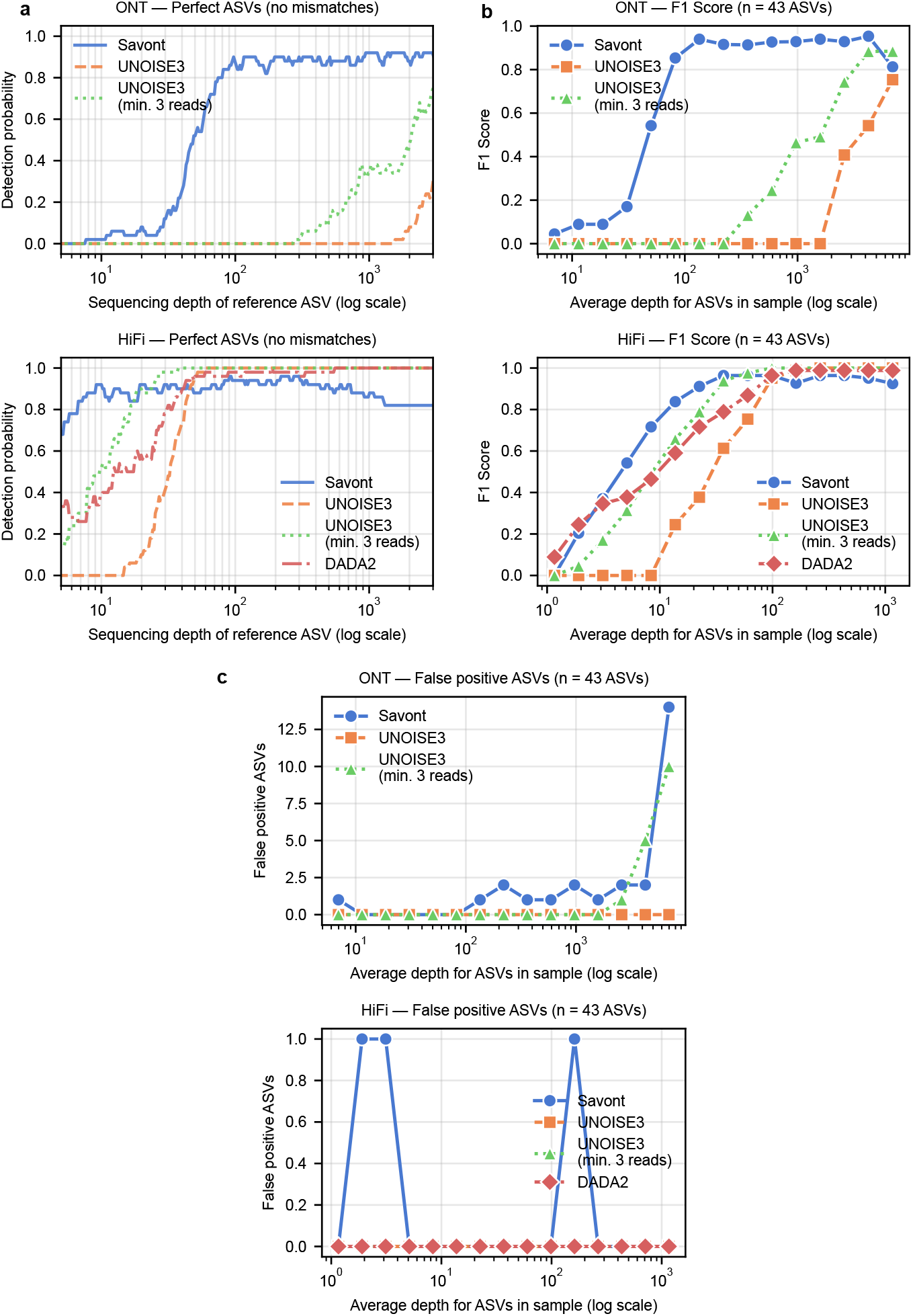
rRNA operon results using full rRNA operon amplicon sequencing datasets from Riisgaard-Jensen et al [14] **a**. Detection probability of ASV as a function of sequencing depth over all pooled samples and using k-nearest neighbor regression with k = 50. **b**. F1 scores for the 43 ASVs per downsampled community. True positives are ASVs with 0 mismatches, and false positives are ASVs with ≥ 1 mismatches to the best reference. **c**. Number of false positive ASVs (number of mismatches ≥ 1) per downsampled community.

**Supplementary Figure 2:**
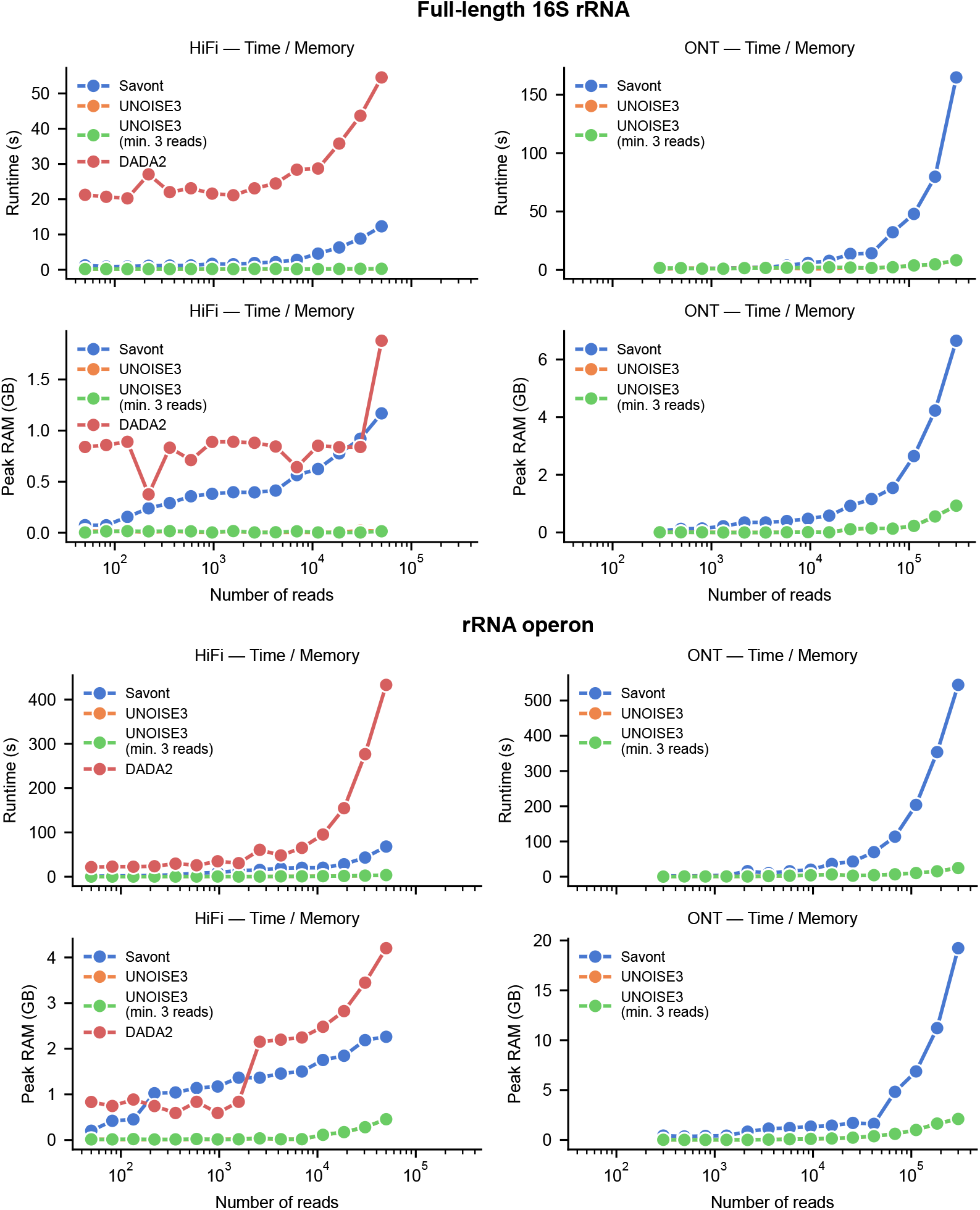
Peak RAM and runtime on the 16S V1-V8 rRNA (top) and rRNA operon (bottom) sequenced downsampled Zymo communities.

**Supplementary Figure 3:**
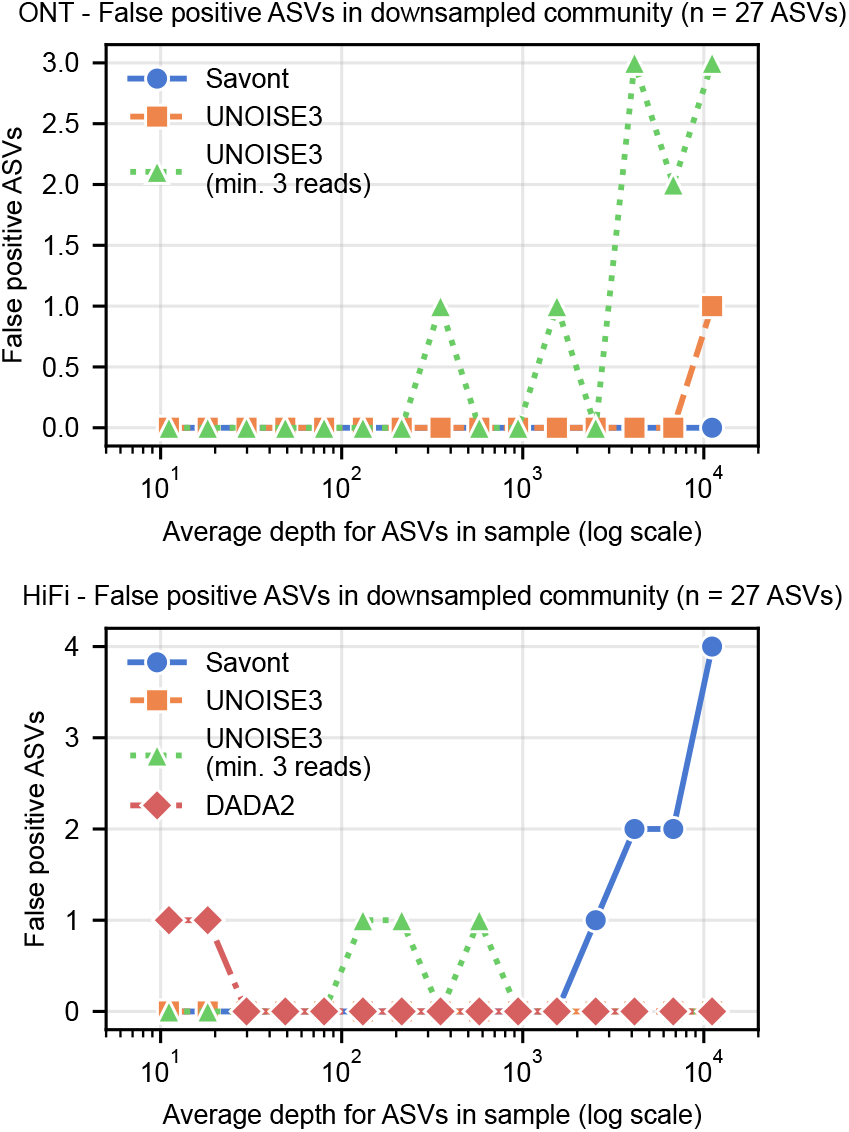
Number of false positive ASVs per downsampled dataset on the Zymo mock community. A false positive was defined as an ASV with *>* 0 mismatches against the 27 reference ASVs.

**Supplementary Figure 4:**
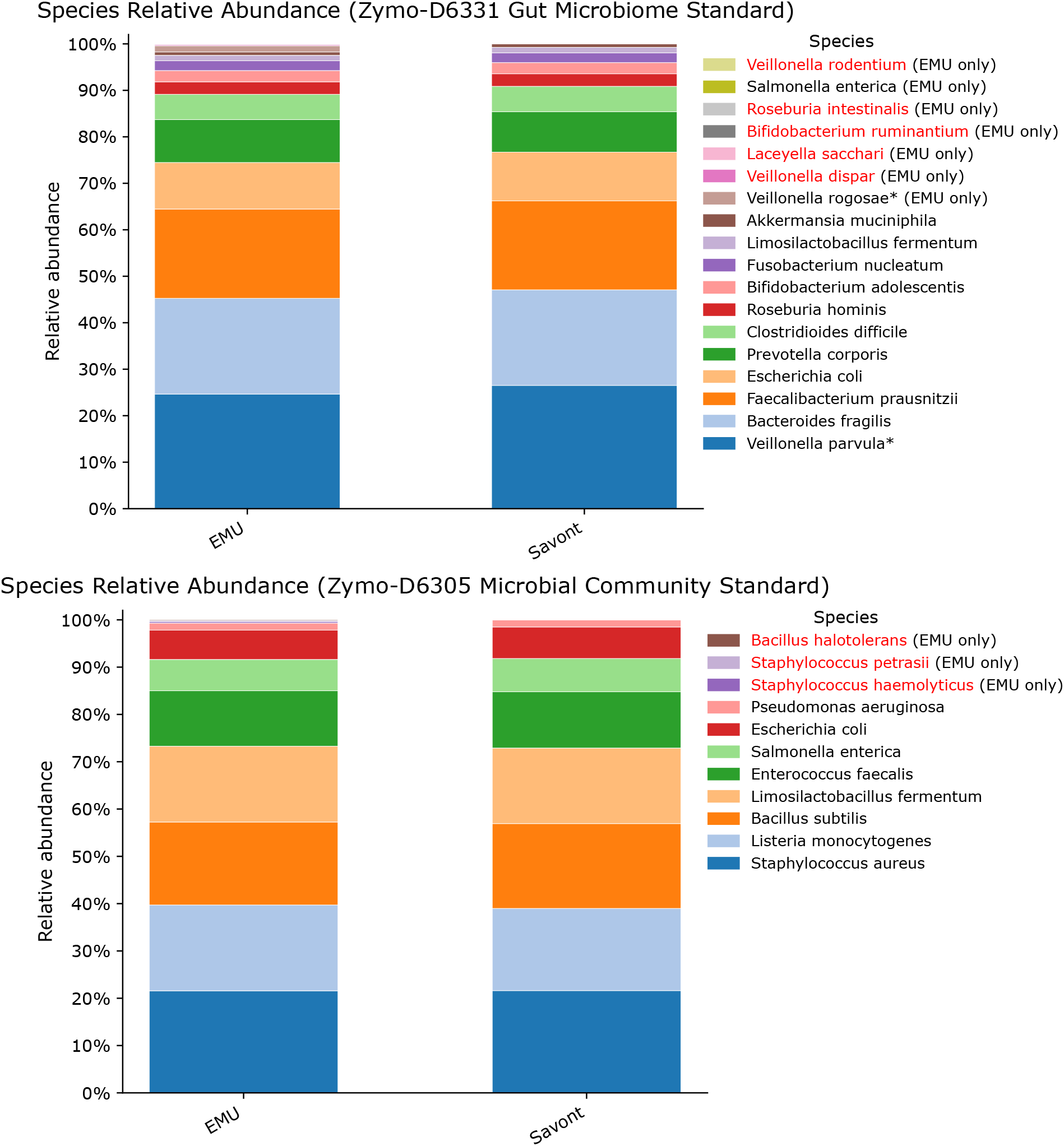
Stacked bar plots for species-level profiling for EMU vs Savont. Top: a Zymo Gut Microbiome Standard full-length 16S rRNA amplicon dataset from ONT (FBB38036 replicate #1). Species not included in the official Zymo specification are highlighted in red. *Veillonella parvula* and *Veillonella rogosae* are marked with an asterisk because while *V. rogosae* is present in the Zymo specification, the true reference ASV (determined by UMIs) has a better alignment to *V. parvula*’s 16S rRNA sequence in EMU’s database. Bottom: the Zymo Microbial Community Standard dataset (technical replicate # 1) from Riisgaard-Jensen et al [14] subsampled to 9,400 reads. Savont was set to classify species-level matches at a relaxed threshold of 98% identity due to the *F. prausnitzii* reference being ≈ 98.8 similar to the ASV.

